# The invasions of *Aedes aegypti* and *Aedes albopictus* in Cyprus: current situation, risk modelling and public health implications for the wider Eastern Mediterranean region

**DOI:** 10.1101/2024.12.17.628941

**Authors:** Cyril Caminade, Marlen I. Vasquez, Herodotos Herodotou, Gregoris Notarides, Costas Pavlou, Filip Fayad, Apostolos Papakonstantinou, Marios Violaris, Dusan Petric, Wadaka Mamai, Adrian M. Tompkins, Jeremy Bouyer

## Abstract

The Asian tiger mosquito, *Aedes albopictus* and the yellow fever mosquito, *Aedes aegypti* have been spreading worldwide and are reshaping the distribution of arboviruses. Both *Aedes* species have recently been observed in densely populated cities of Cyprus, a touristic island that is a historic bridge between Europe and Asia. Given the high public health stakes for Cyprus and the wider East Mediterranean region, the objectives of this study are three-fold. First, we present a novel delimitation strategy using spatially dense networks of ovitraps deployed in 500×500m cells in Limassol and Larnaca following the detection of *Aedes* species. Second, we use a dynamical vector model to estimate the potential of both species to spread further over Cyprus. Finally, we employ a basic reproduction number (R_0_) model to assess the potential transmission risk of arboviruses for the wider East Mediterranean region. Our results underline our delimitation strategy’s usefulness in delineating *Ae. albopictus* populations in Limassol and indicate the need for increased surveillance efforts for *Ae. aegypti* in Larnaca. Our vector model reveals that cities such as Nicosia, Paphos and Ayia Napa are climatically suitable for the establishment of both *Ae. aegypti* and *Ae. albopictus*. Finally, the R_0_ model captures historical hotspots of dengue transmission over the East Mediterranean region, with large R_0_ values simulated over Cyprus, Greece, Turkey, southern Italy and southern Spain. We recommend stringent vector surveillance at entry points in Greece and a rapid elimination in Cyprus to prevent the return of *Ae. aegypti* to the European continent.

## Introduction

*Aedes* mosquitoes are competent vectors of numerous pathogens, such as *Dirofilaria immitis*, the dog heartworm, and important arboviruses such as dengue, yellow fever, chikungunya and Zika. These arboviruses are a major threat to human life, especially in tropical and subtropical regions. *Aedes aegypti* (Linnaeus, 1762) and *Ae. albopictus* (Skuse, 1895) are the main vectors of important arboviruses impacting public health. Both species have rapidly spread worldwide due to globalization and the movement of goods (Cuthbert et al., 2023), and their establishment was favored by more suitable climatic conditions (Caminade et al., 2012).

The Asian tiger mosquito, *Ae. albopictus*, was introduced into Albania from Asia during the late 1970s, very likely by exports of used tyres and/or lucky bamboo plants (Cuthbert et al., 2023). This species then rapidly spread to Italy, France, Montenegro, Greece and Switzerland and is now broadly established over southern and central Europe (ECDC, 2024a). Its recent spread in Europe was favoured by ground vehicles following the motorway network, trains and boats (Swan et al., 2022) as well as increasingly suitable climatic conditions (Caminade et al., 2012). The yellow fever mosquito, *Ae. aegypti*, used to be historically widespread over the Mediterranean and Adriatic coasts of southern Europe and was responsible for the largest historical dengue epidemic that occurred in Athens in 1927-28 (Wint et al., 2022). *Aedes aegypti* disappeared from southern European countries following large DDT malaria vector control campaigns post World War II, management of urban water collection and possibly a lower potential to establish itself due to cold winter conditions (Christophers, 1960). Nowadays, *Ae. aegypti* is mostly restricted to the eastern coasts of the Black sea and north-eastern Turkey (ECDC, 2024a). Both species are daily biting nuisances, complementing to the risk of pathogen transmission by native nocturnal vectors. *Aedes albopictus* has been increasingly responsible for autochthonous dengue and chikungunya cases in southern France, Italy, Croatia and Spain over the past few years (ECDC, 2024b). Notably, 65 and 82 autochthonous cases of dengue were respectively reported in France in 2022 and in Italy in 2023. *Ae. aegypti* was also responsible for a large dengue outbreak in Madeira in 2012 with more than 2000 local cases and 81 exported cases to mainland EU (Lourenço & Recker, 2014).

In Cyprus, *Ae. aegypti* was found in historical surveys before 1934 (Violaris et al., 2009), then disappeared in the 1950s. Its recent re-introduction and detection in the vicinity of the international airport of Larnaca, is worrying given its high competence to transmit yellow fever and dengue. *Aedes albopictus* was also recently detected in the old town area of Limassol (UNDP, 2024, Vasquez et al., 2023). These recent (re)introductions are extremely worrying, given probable suitable climatic conditions, a very well-connected network of motorways and the up to 4 million tourists visiting Cyprus each year.

Given such high stakes for public health in Cyprus and the wider East Mediterranean region, the objectives of this study are three-fold. First, we present a novel trapping strategy that was used in Cyprus from October 2022 until June 2023 to delimitate the spread of both *Aedes* species at high spatial resolution. Second, we utilize a climate driven mathematical model to map potential invasion hotspots as well as seasonal activity of both *Aedes* species in Cyprus. Finally, we use a generic basic reproduction number (R_0_) model for arboviruses in order to provide recommendations to public health experts in Cyprus and for the wider Eastern Mediterranean region.

## Methods

### Delimitation strategy

A detailed delimitation strategy was implemented to understand the spread of both *Ae. Aegypti* in Larnaca (Vasquez et al., 2023) and *Ae. albopictus* in Limassol following their detection (Fig. S1). A 500m x 500m grid was designed for each area under study, and a unique ID number was given to each cell. “Infested cells” were defined if invasive *Aedes* eggs or adults were collected. We aimed to survey invasive *Aedes* within a 5km radius from each infested cell. Sampling priority was given to cells adjacent to infested cells, then cells 1000m from infested cells and so on. Each cell was surveyed at least 3 times in the case of absence of capture, using 10 ovitraps set for at least two weeks or by conducting Human Landing Catch (HLC) in 5 sites within the cell for 5 minutes. The trapping methods were alternated so as to survey each cell at least twice with ovitraps. The use of ovitrap aimed to overcome any bias that may arise from the HLC method with regard to various host preferences and varying levels of collector expertise.

A “traffic light” system was used to prioritize the areas to survey. If any specimens of invasive *Aedes (Stegomyia)* species were collected, the cells were considered as “positives” (red coloured). “Suspicious cells” (yellow coloured) were considered based on citizen science data or for cells adjacent to positive cells. “Cells under investigation” (grey coloured) were considered for all cells with less than 3 surveys. “Provisionally free” were cells with no collection following 3 surveys separated by 10 weeks (green coloured). The highest priority was given to yellow cells during the delimitation survey.

A data collection and management system was developed using ArcGIS Survey123, and maps were produced weekly to guide the field entomologists to the survey areas. Ten teams of two health workers from the Ministry of Health, the local health authority (i.e. municipality, community, UK Sovereign Base Area of Akrotiri) and CUT researchers were deployed in each area for ovitrap placement and collection, whereas HLC was carried out by two CUT researchers. Following the training of the health workers, the delimitation area increased from 10 to 100 cells/week in each area within two months in Larnaca and within one month in Limassol area.

### Entomological data

The “delimitation data” is based on a spatially dense network of ovitraps placed in Limassol and Larnaca from October 2022 until June 2023. The identification of eggs was inferred by the invasive *Aedes* adult catches using HLC during the same period. Ovitrap data was converted to the number of eggs per trap per day (or per week) to allow comparability across study sites.

Long-term historical occurrence data for *Ae. aegypti* was derived from the recent data compiled and curated by Schaffner, 2022. Occurrence data for *Ae. albopictus* was derived from the Global Biodiversity Information Facility (*Aedes albopictus*, GBIF, 2023) for comparison purposes.

### Climate and population density data

The European Meteorological Observations 1arcmin (EMO-1) climate data (Gomes et al., 2020) was utilized to drive the VECTRI model (VECtor borne disease community model of ICTP, TRIeste). The EMO-1 dataset has a spatial resolution of 1arcmin x 1arcmin (approx. 1.5km x 1.5km) and is available at a daily time step over the period 1990-2022. We used rainfall (in mm) and mean temperature (in °C) to drive the *Ae. aegypti* and *Ae. albopictus* VECTRI models. EMO-1 is routinely used by the European Joint Research Centre (JRC) to provide flood risk assessments. It is noteworthy that other simulations using the CHELSA climate daily data (1km x 1km spatial resolution for the period 1985-2005; Karger et al., 2020) to drive the VECTRI model were carried out. As CHELSA results were consistent with EMO-1 (not shown), we focus on the EMO-1 dataset because this dataset was available until the end of 2022. Population density data (per km^2^) is based on the GPwv4 dataset (Doxsey-Whitfield et al., 2015). This dataset was used as an input to the VECTRI model and to highlight spatial overlaps between large human density and simulated abundance hotspots.

### Aedes vector simulations

VECTRI is a mathematical malaria model that includes the effects of precipitation, temperature, population immunity, hydrology (using a simple pool model), and human population density. The model was originally developed for *Anopheles gambiae* and *Plasmodium falciparum* (Tompkins & Ermert, 2013) but it was recently adapted to *Ae. albopictus* and *Ae. aegypti*. VECTRI dynamically models the different life cycles of the mosquito vector and their daily dependency on rainfall and temperature. VECTRI also takes into account hydrological surface conditions and human population density. As both *Aedes* mosquito species thrive in urban environments, due to the availability of person-made water containers, their dependency on rainfall was lowered (wperm value was lowered to 0.5 x 10^-6^). Adult mortality schemes dependency to temperature were derived from Metelmann et al., 2019 for *Ae. albopictus* and Brady et al. 2013 for *Ae. aegypti*. The updated version of the VECTRI model for *Aedes* species is available on Gitlab at [https://gitlab.com/tompkins/vectri/-/tree/aedes?ref_type=heads]. Model outputs for *Ae. aegypti* and *Ae. albopictus* are available on the Open Science Framework platform at https://osf.io/7h5du/.

### Basic reproduction number (R_0_) model to estimate the risk of arbovirus transmission

We utilised the two vectors - one host R_0_ model that was developed to assess the risk of Zika virus transmission at global scale (Caminade et al., 2017). This basic reproduction number model considers the two *Aedes* vectors under scope, namely *Ae. aegypti* and *Ae. albopictus*. Even though this model was originally developed to assess the risk of Zika transmission, the original parameter setting was mostly based on dengue parameters. This R_0_ model has subsequently been used to estimate the generic risk of *Aedes*-borne disease transmission by Muñoz et al. 2020. The model was driven by observed precipitation and temperature data at 0.5°x0.5° for the period 1948-2015. Model outputs are available at https://osf.io/ubwya/

## Results

The delimitation ovitrap data that was collected in Cyprus is shown in Figure 1. Positive *Ae. aegypti* traps are primarily located in the Larnaca area, in the vicinity of the international airport and in the southern part of the city centre (Fig 1a). *Aedes aegypti* was not collected in any other location apart from Larnaca, indicating the current geographical restriction of this species. Positive *Ae. albopictus* traps are shown over Limassol (Fig 1b). *Aedes albopictus* was primarily found by the old port, the marina and the eastern coastal part of the city. It is noteworthy that *Ae. albopictus* was detected in an eastern area of Cyprus (Agios Tychon) where another member of genus Stegomyia, *Aedes cretinus*, which female closely resemble *Ae. albopictus*, is known to be present (not shown). Positive *Ae. albopictus* traps were also collected in Paphos and Nicosia in September 2023 by local authorities and verified by CUT researchers (not shown). Delimitation activities in these areas are taking place in 2024 (not shown).

**Fig. 1:**
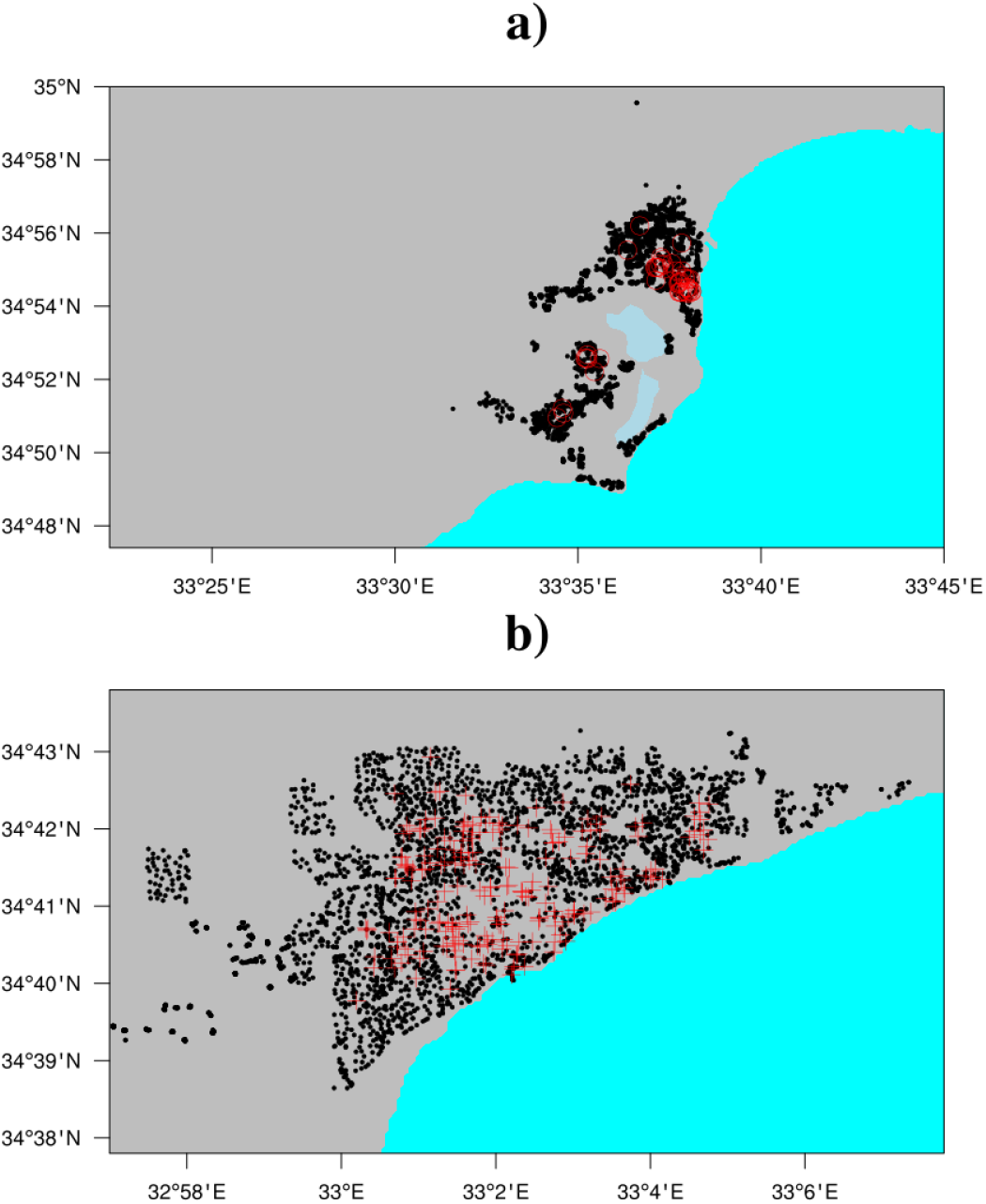
Ovitrap data collected between Oct 2022 until June 2023 over a) the area of Larnaca (Ae. aegypti) and b) the area of Limassol (Ae. albopictus). Negative traps are depicted by black dots, positive Ae. aegypti traps are denoted by red circles and positive Ae. albopictus traps are indicated by red crosses.

In order to explore the delimitation ovitrap data into greater details, the number of positive ovitraps per 500×500m cell is depicted for *Ae. aegypti* in Larnaca (Fig 2a-b) and *Ae. albopictus* in Limassol (Fig 2c-d). Most traps were negatives. Most positive cells had only one positive trap and positive traps for all sampled cells showed a negative binomial distribution (Fig 2b-d). The spatial coverage and number of ovitraps deployed in Limassol perfectly delineated *Ae. albopictus* positives with respect to negatives (depicted by the white area on Fig 2c). The number of ovitraps deployed in the Larnaca area and the associated spatial coverage for *Ae. aegypti* was lower (Fig 2a).

**Fig. 2:**
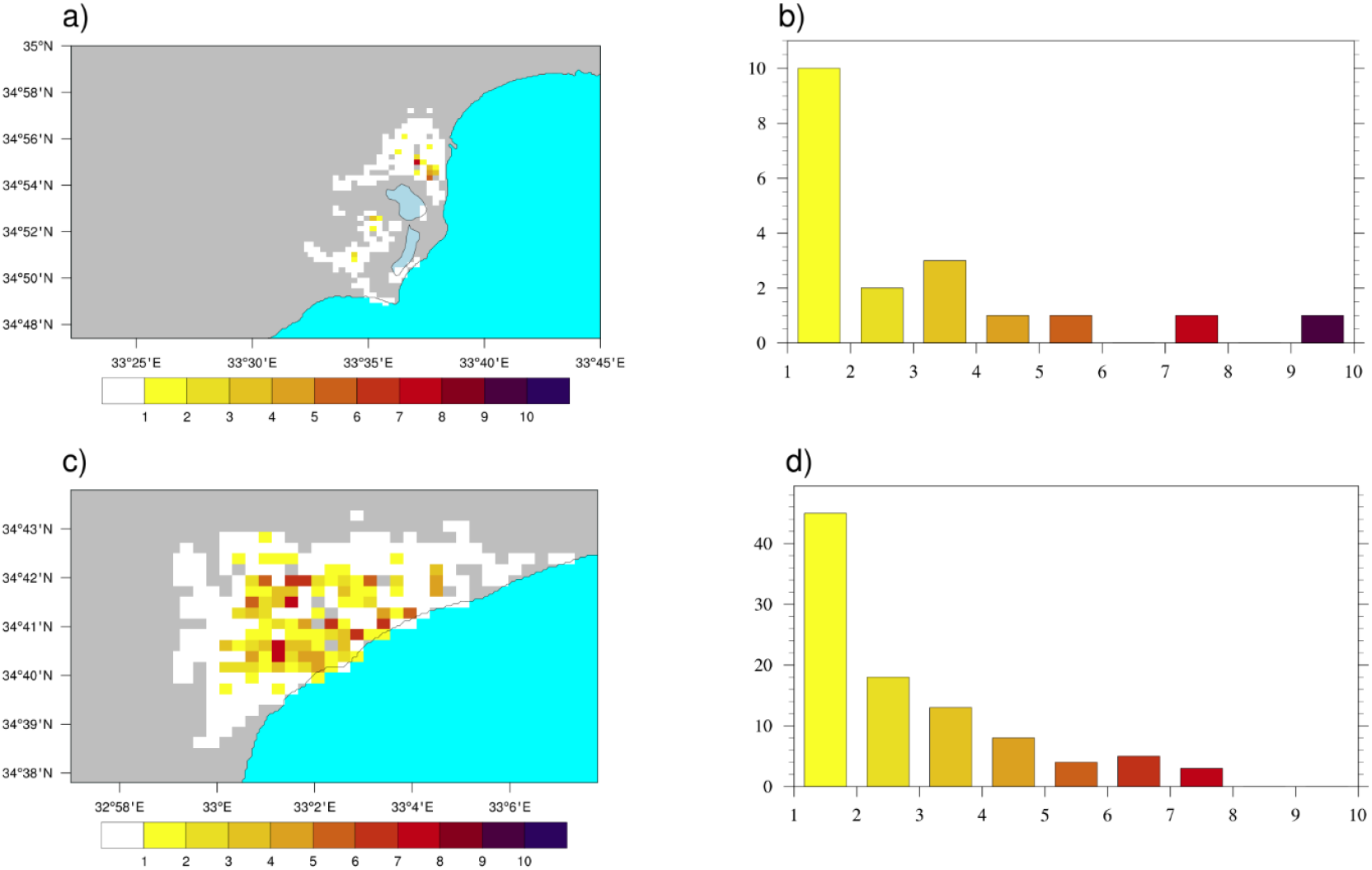
Number of positive traps per 500×500m cell for (a) the area of Larnaca (Ae. aegypti) and (c) the area of Limassol (Ae. albopictus). Negative traps are depicted in white on the maps. Histogram of the number of positive traps for all cells in b) the area of Larnaca and d) the area of Limassol.

Abundances of *Ae. aegypti* eggs in Larnaca and *Ae. albopictus* eggs in Limassol are respectively shown on Fig. 3a and Fig. 3b. Largest abundances of *Ae. aegypti* were observed in the southern part of Larnaca city centre, with more moderate abundance found in the vicinity of Dromoloxia-Meneou, south of Larnaca international airport (Fig. 3a). Largest *Ae. albopictus* egg abundance was found in the old port, Neapolis and the marina area of Limassol as well as around the Kato Polemidia area (Fig. 3b). Time animations highlight rapid spatial jumps of *Ae. albopictus* from the old city to the eastern coasts of Limassol (not shown).

**Fig. 3:**
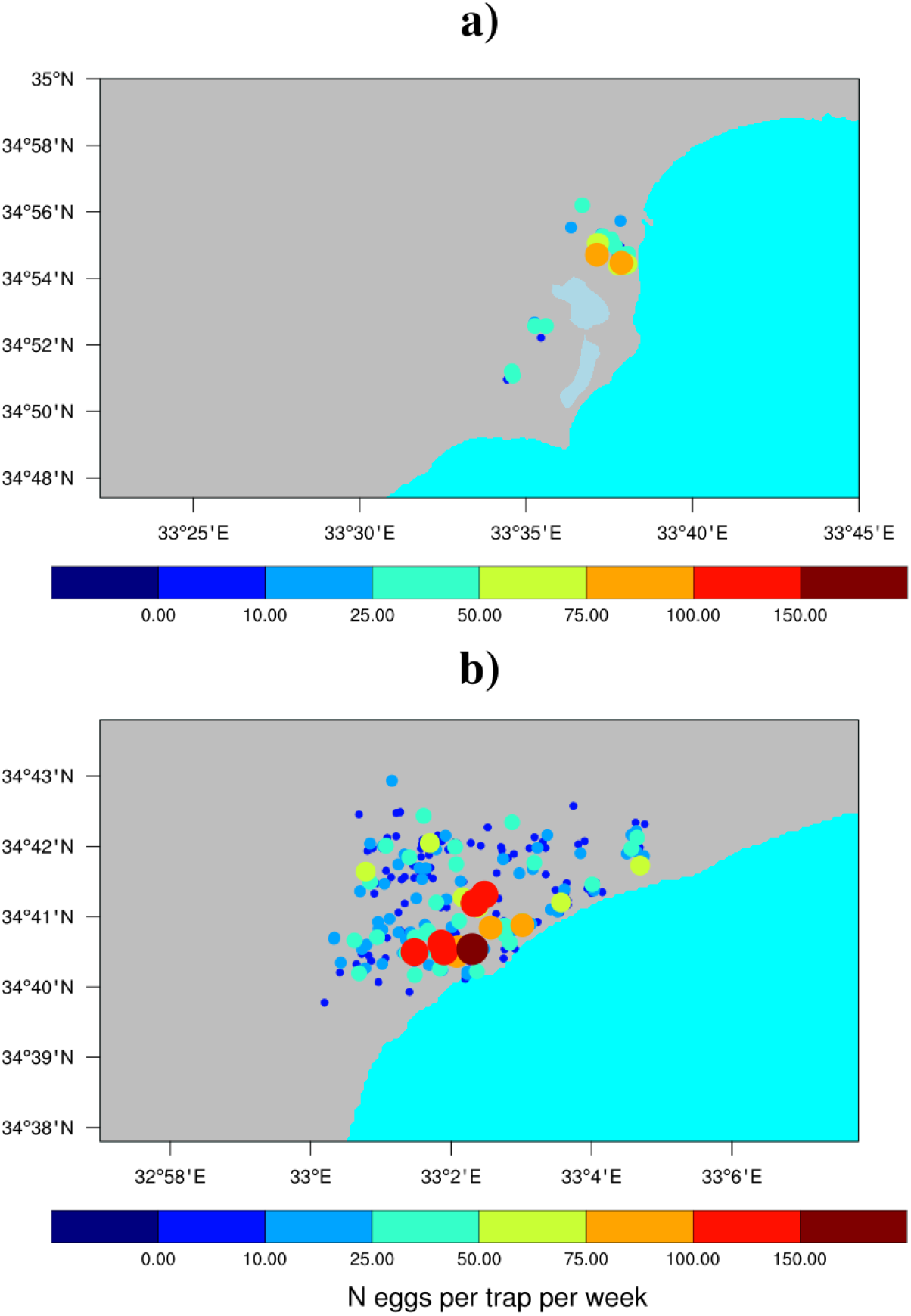
Egg abundance (per trap per week) for a) Ae. aegypti in Larnaca area and b) Ae. albopictus in Limassol area. Only positive traps are shown, see Fig. 1 for negatives.

The VECTRI model was driven by observed daily rainfall and temperature data at high spatial resolution (about 1.5×1.5km) over Cyprus for the period 1990-2022. Heat maps for both *Ae. albopictus* and *Ae. aegypti* are shown in Figure 4. Cold spots are shown over the altitude regions of Cyprus, namely over the western Troodos Mountains and over the northern Kyrenia Mountains. Hotspots are highlighted over densely populated cities of Cyprus such as Limassol, Paphos, Larnaca (Fig. 4 and Fig. S2) and, to a lesser extent, Ayia Napa. Large values are also shown over Nicosia, Morphou, over the eastern and northern coasts of Cyprus. Simulated larval density values increase significantly when the model is driven by recent climate conditions (2010-22 vs 1990-2009; Fig. 4b-d vs Fig. 4a-c). Interestingly, the low-altitude regions appear more at risk of establishment during the 2010s. Both mosquito species could theoretically spread to densely populated areas in Cyprus, and notably the capital Nicosia.

**Fig. 4:**
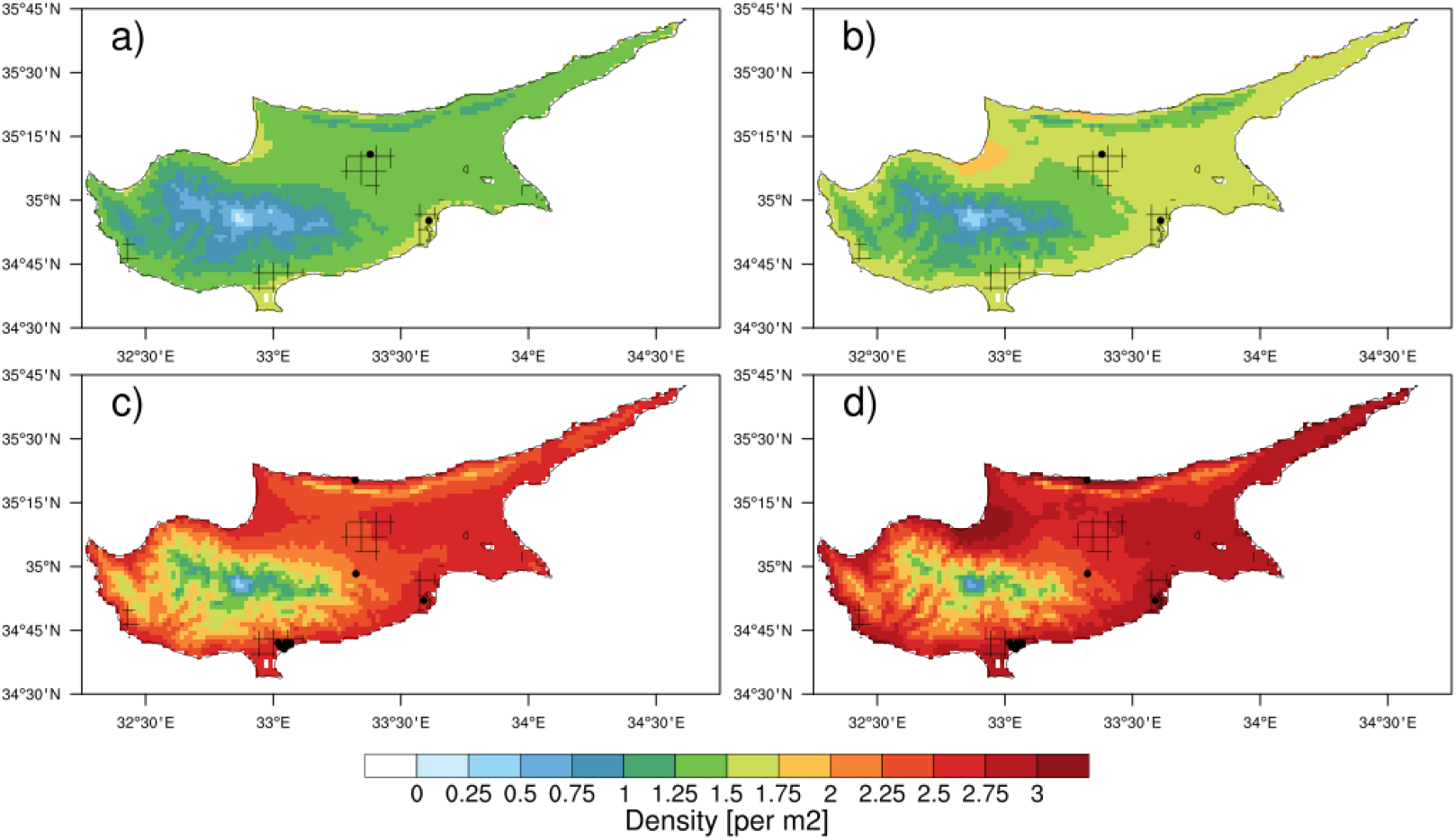
Average larval density (per m^2^) as simulated by the VECTRI model for Ae. aegypti (a, b) and Ae. albopictus (c, d) for 1990-2009 (a, c) and 2010-22 (b, d). Historical occurrences of Ae. aegypti (Schaffner et al., 2022) and Ae. albopictus (GBIF, 2024) are depicted by black dots. Human population density above 100 people per km^2^ is depicted by the hatched area.

A peak in *Ae. aegypti* eggs was observed from late October until mid November 2022 in Larnaca (Fig. 5a). *Aedes albopictus* eggs were found in the Limassol area from late October until mid-December 2022 (Fig. 5b). Almost no eggs were trapped during Jan-Feb 2023. Then, first eggs were collected during mid-March 2023 and continued until the end of the delimitation period in June 2023 (Fig. 5b). Ovitrap data reveals that *Ae. albopictus* starts being active in late March-early April in Limassol and its abundance increases during the late spring and early summer months (Fig. 5b). Simulated larval density for *Ae. albopictus* tends to be twice larger compared to *Ae. aegypti* (Fig. 4). The mean seasonal cycle maps of *Ae. aegypti* and *Ae. albopictus* are respectively shown in Fig. S3 and Fig. S4. Simulated larval density for *Ae. albopictus* are consistently larger compared to *Ae. aegypti* for all months. In our simulations, both mosquito species start being active in April and are still active in December over the coastal and eastern part of Cyprus. Large larval densities are simulated from May until October. The minimum is simulated for both species during boreal winter from January until March. Simulated values are systematically lower over altitude regions, namely over the Troodos and Kyrenia mountains. Simulated values are large over Nicosia in June and September for *Ae. albopictus*.

**Fig. 5:**
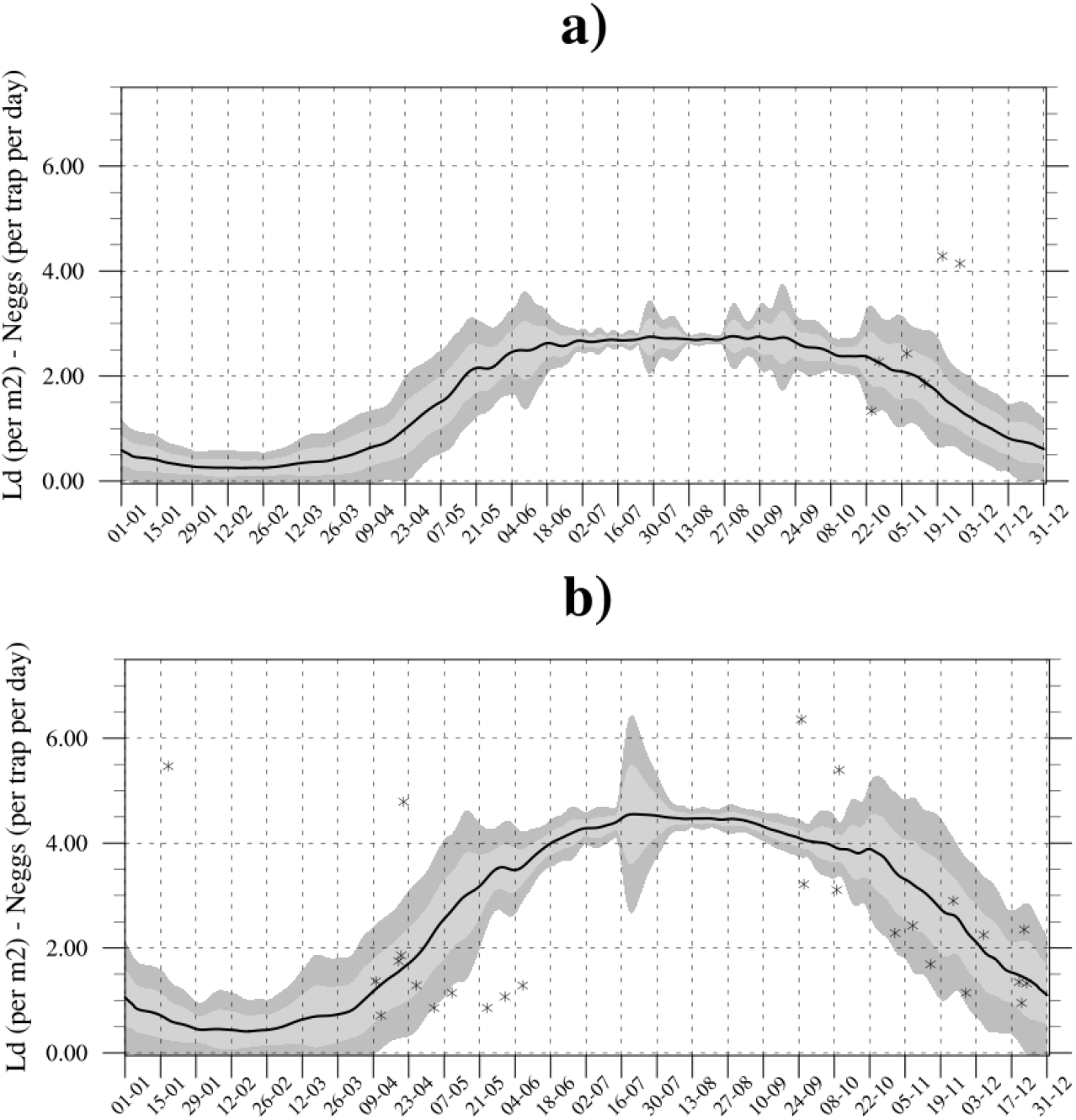
Mean seasonal cycle of simulated larval density for a) Ae. aegypti in Larnaca and b) Ae. albopictus in Limassol. The mean for 1990-2022 is depicted by the black solid line. The light-gray and gray envelope respectively show one and two standard deviations around the mean calculated for all years. Observed abundance of eggs (median number of eggs per trap per day) are depicted by black crosses for the period Oct 2022 – June 2023.

Overall, large larval density values are shown over densely populated urban centres and over the eastern part of Cyprus during the summer months. It is noteworthy that another set of VECTRI simulations driven by maximum temperature was carried out (not shown). These simulations reveal that higher daily temperature conditions in July-August decrease significantly simulated larval density for both species during mid-summer over Larnaca, Limassol and Nicosia, resulting in a bi-modal seasonal cycle shape (with peaks in boreal spring and autumn, see Fig. S5). In 2020, climate suitability for both *Aedes* species significantly decreased in Nicosia during summer. Such decrease can directly be related to heatwave conditions that have detrimental effects on simulated abundance (not shown).

The presence of *Aedes* vectors that are competent to transmit arboviruses is an important factor, however other drivers need to be considered to estimate potential disease transmission risk. We estimate the risk of arbovirus disease transmission by both *Ae. aegypti* and *Ae. albopictus* using a basic reproduction number (R_0_) model for the East Mediterranean region and the period 1980-2010 (Fig. 6). Large R_0_ values (> 1) are simulated over the Attica region and Crete in Greece, Cyprus, the western and southern coasts of Turkey, southern Italy and the coasts of Tunisia and Algeria. These hotpots are consistent with historical dengue epidemics reported in Cyprus in 1861, 1888-89, 1913 and 1928; in Crete in 1881; over western Greece in 1889, 1895-97,1910, 1927-28, 1929-33; in Napoli in 1889-90, and in Turkey in 1889-90, 1899, 1916, 1927-28 and 1945 (Schaffner & Mathis, 2014). The model also simulates large R_0_ values over Andalucía in Spain (not shown) that experienced dengue epidemics in 1784-88-93, 1863-67, 1887 and 1928.

**Fig. 6:**
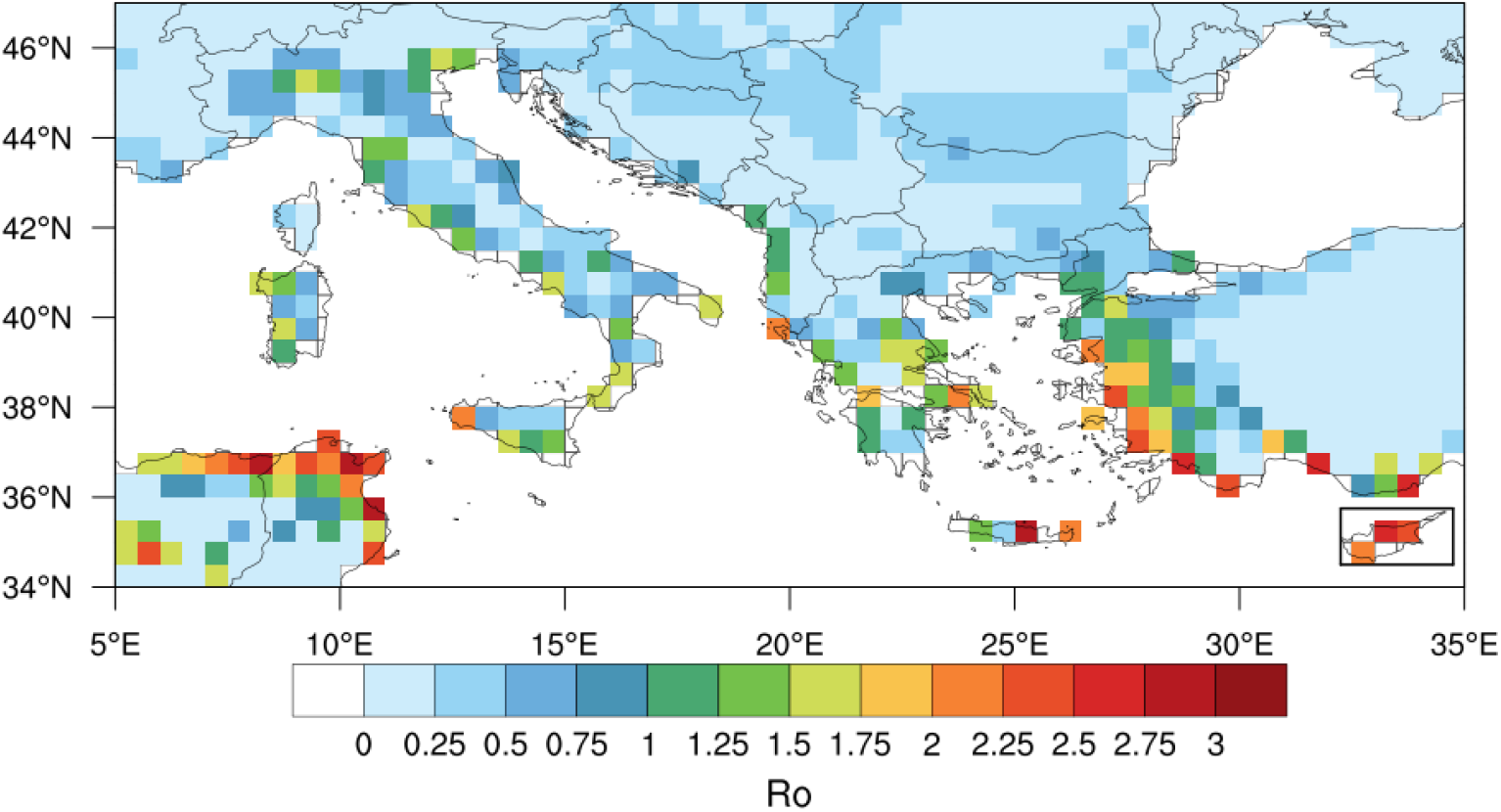
Annual mean basic reproduction number (R_0_) for the risk of arbovirus transmission by *Ae. aegypti & Ae. albopictus* over the East Mediterranean region (1980-2010). Cyprus is highlighted by the black box.

## Discussion

A novel delimitation technique applied to *Aedes* mosquitoes highlighted that *Ae. albopictus* is already well established in Limassol, but also Paphos and Nicosia in Cyprus (Vasquez et al., 2023). Eggs were found in Limassol from mid-March until late December, with a peak recorded in September. *Aedes aegypti* has been primarily found in Larnaca, in the Dromoloxia-Meneou area, close to the international airport and in the southern part of the city centre. *Aedes aegypti* eggs were found in Larnaca area, with a peak observed in mid-November. The VECTRI model reproduces realistic seasonal activity of both *Ae. aegypti* and *Ae. albopictus*. On average, the simulated length of the activity season for *Ae. albopictus* is slightly longer (early April until late-December) with respect to *Ae. aegypti (*late April until early-December). This finding is consistent with the fact that strains of *Ae. albopictus*, with diapausing eggs, are well adapted to temperate climate conditions with a large potential to overwinter, while *Ae. aegypti* tends to thrive in semi-tropical climes; its eggs are less resilient to cold winter conditions (Kramer et al., 2021).

Simulations highlight that coastal areas of Cyprus are climatically suitable for the establishment of both *Aedes* species in densely populated cities, such as Paphos, Limassol, Larnaca, Ayia Napa, and Nicosia. Large larval density values are also simulated near Morphou, over the eastern part and the northern coasts of Cyprus where *Ae. albopictus* was reported to be present by a citizen science project (*Aedes albopictus*, GBIF, 2023). It is noteworthy that *Ae. aegypti* is primarily anthropophilic, and should therefore be found in densely populated urban centres. *Aedes albopictus* is a more opportunistic feeder (birds, rodents and other mammals) and consequently can survive in peri-urban areas but it is also extremely well adapted to urban settings. Recent climate change conditions during the 2010s favoured an increase in simulated larval density, particularly over the coastal regions and over the fringes of the Troodos and Kyrenia mountains. This feature is consistent with other studies showing the spread of *Ae. albopictus* at higher elevations in Europe (Romiti et al., 2022; Tisseuil et al., 2018).

Our modelling framework does not consider the movement of hosts or vectors over long distances, as well as several important socio-economic factors (vulnerability factors, human behaviour, etc.) might impact vector distribution. Mosquitoes can also adapt, evolve and compete. Such evolutionary aspects were not included in our study. We produced high spatial resolution mosquito density estimates (1.5 x 1.5 km) but we did not consider micro-climatic conditions (shaded areas) or land types.

Based on our findings, stringent and continuous surveillance effort should be developed and intensified in Larnaca but also Nicosia, Ayia Napa, Morphou and over the northern coast and eastern part of Cyprus. Stringent surveillance of imported dengue, Zika and chikungunya cases in spring-summer in portal cities and at Larnaca international airport and a strong control or elimination response should be conducted. A published epidemiological study found that the introduction of dengue cases to Madeira in 2012 occurred around a month before the first autochthonous cases were reported, during the maximum influx of airline travellers, and that declining temperature conditions in autumn were critical in limiting the lifespan of the dengue outbreak in early December 2012 (Lourenço & Recker, 2014). Importantly, the presence of *Ae. aegypti* was first detected in the Funchal area in 2005, seven years before the dengue outbreak occurred in Madeira (Gonçalves et al., 2008). Given that Greece is the main trading partner of Food products into Cyprus (World Bank, 2021), stringent surveillance should be conducted in containers transiting between the main harbours of both countries.

Finally, as a wake-up call, the largest dengue epidemics occurred in Greece, Cyprus, Turkey, southern Italy and southern Spain during the first half of the 20^th^ century, in a pre-Anthropocene era. These epidemics were very likely caused by the yellow fever mosquito *Ae. aegypti*, before its disappearance post-WWII. Hence it is of paramount importance to limit the spread of this species to the wider Eastern Mediterranean region. A contingency strategy has thus been proposed by the IAEA to attempt to eliminate *Ae. aegypti* from Cyprus using an integrated vector management strategy including a Sterile Insect Technique (SIT) component, and a pilot trial SIT trial has been initiated in Larnaca in 2023.

## Funding

CC, AMT and JB acknowledge internal funding from the ICTP-IAEA project “Using the VECTRI mosquito model with machine learning to derive optimal strategies for the Sterile Insect Technique (SIT)”. This study was supported by the national IAEA TC project CYP5020 “Developing a national rapid response strategy for the prevention of the establishment of the Asian tiger mosquito” (equipment and consumables).

## Acknowledgements

The delimitation strategy was supported by health workers of the MPHS and CUT researchers. The National Surveillance program was supported by CUT researchers.

## Data

The entomological data is available under request to Marlen Vasquez. VECTRI model outputs are available at [https://osf.io/7h5du/]. R_0_ model outputs are available at [https://osf.io/ubwya/].

## Authors’ contributions

GN, MIV, HH, CP, MV and JB supervised field campaigns. DP, JB and MIV conceptualized the principles of the delimitation strategy. FY and AP developed and maintained the GIS database, AMT and CC adapted the VECTRI model, MIV and CC curated the database, CC carried out the analysis and wrote the manuscript with inputs from all co-authors.

## Notes

### Competing Interest Statement

The authors have declared no competing interest.

## References

Aedes albopictus (Skuse, 1895) in GBIF Secretariat (2023). GBIF Backbone Taxonomy. Checklist dataset 10.15468/39omei accessed via GBIF.org on 2024-01-25.

Brady, O.J. et al. (2013). Modelling adult Aedes aegypti and Aedes albopictus survival at different temperatures in laboratory and field settings. Parasites Vectors 6, 351

Caminade C., J.M. Medlock, S. leach, K.M. McIntyre, M. Baylis, A.P. Morse (2012). Suitability of European climate for the Asian tiger mosquito Aedes Albopictus: recent trends and future scenario. Journal of the Royal Society Interface. 9(75): 2708–2717

Caminade C., J. Turner, S. Metelmann, J.C. Hesson, M.S.C. Blagrove, T. Solomon, A.P. Morse, M. Baylis (2017). Global risk model for vector-borne transmission of Zika virus reveals the role of El Nino 2015. PNAS, 114(1): 119–124.

Christopher S. Rickard (Sir) (1960). The Yellow Fever mosquito. Its life history, bionomics and structure. Cambridge University Press. New York, 1960, 739 pp.

Cuthbert, R.N., Darriet, F., Chabrerie, O. et al. (2023). Invasive hematophagous arthropods and associated diseases in a changing world. Parasites Vectors 16, 291 (2023).

Doxsey-Whitfield E. et al. (2015) Taking Advantage of the Improved Availability of Census Data: A First Look at the Gridded Population of the World, Version 4, Papers in Applied Geography, 1:3, 226–234

European Centre for Disease Prevention and Control and European Food Safety Authority. Mosquito maps [internet]. Stockholm: ECDC; (2024a). Available from: https://ecdc.europa.eu/en/disease-vectors/surveillance-and-disease-data/mosquito-maps

European Centre for Disease Prevention and Control. Autochthonous vectorial transmission of dengue virus in mainland EU/EEA, 2010-present. Stockholm: ECDC; (2024b). Available from: https://www.ecdc.europa.eu/en/all-topics-z/dengue/surveillance-and-disease-data/autochthonous-transmission-dengue-virus-eueea

Gomes, Goncalo et al. (2020): EMO: A high-resolution multi-variable gridded meteorological data set for Europe. European Commission, Joint Research Centre (JRC) [Dataset] doi: 10.2905/0BD84BE4-CEC8-4180-97A6-8B3ADAAC4D26 PID: http://data.europa.eu/89h/0bd84be4-cec8-4180-97a6-8b3adaac4d26

Gonçalves, Y. et al. (2008). On the presence of Aedes (Stegomyia) aegypti Linnaeus, 1762 (Insecta, Diptera, Culicidae) in the island of Madeira (Portugal). Bol. Mus. Mun. Funchal 58, 53–59.

Karger, D.N., Dabaghchian, B., Lange, S., Thuiller, W., Zimmermann, N.E., Graham, C.H. (2020): High resolution climate data for Europe. EnviDat. 10.16904/envidat.150.

Kramer IM, Pfeiffer M, Steffens O, Schneider F, Gerger V, Phuyal P, Braun M, Magdeburg A, Ahrens B, Groneberg DA, Kuch U, Dhimal M, Müller R. (2021). The ecophysiological plasticity of Aedes aegypti and Aedes albopictus concerning overwintering in cooler ecoregions is driven by local climate and acclimation capacity. Sci. Total Environ., 778: 146128.

Lourenço J, Recker M (2014). The 2012 Madeira Dengue Outbreak: Epidemiological Determinants and Future Epidemic Potential. PLoS Negl. Trop. Dis. 8(8): e3083.

Metelmann S. et al. (2019). The UK’s suitability for Aedes albopictus in current and future climates. Journal of the Royal Society Interface 16: 20180761.

Muñoz, Á.G., Chourio, X., Rivière-Cinnamond, A. et al. (2020). AeDES: a next-generation monitoring and forecasting system for environmental suitability of Aedes-borne disease transmission. Sci Rep 10, 12640 (2020).

Romiti, F., Casini, R., Magliano, A. et al. (2022). Aedes albopictus abundance and phenology along an altitudinal gradient in Lazio region (central Italy). Parasites Vectors 15, 92.

Schaffner F, Mathis A. (2014). Dengue and dengue vectors in the WHO European region: past, present, and scenarios for the future. Lancet Infect Dis. 12:1271–80.

Schaffner, Francis (2022). Historical and modern distribution data of the yellow fever mosquito Aedes aegypti in the western Palaearctic region. Figshare dataset. 10.6084/m9.figshare.20343570.v1

Swan T, Russell TL, Staunton KM, Field MA, Ritchie SA, Burkot TR. (2022). A literature review of dispersal pathways of Aedes albopictus across different spatial scales: implications for vector surveillance. Parasite & Vectors, 15(1):303.

Tisseuil C, Velo E, Bino S, Kadriaj P, Mersini K, Shukullari A, Simaku A, Rogozi E, Caputo B, Ducheyne E, Della Torre A, Reiter P, Gilbert M (2018). Forecasting the spatial and seasonal dynamic of Aedes albopictus oviposition activity in Albania and Balkan countries. PLoS Negl Trop Dis. 12(2)

Tompkins, A.M., Ermert, V. (2013). A regional-scale, high-resolution dynamical malaria model that accounts for population density, climate and surface hydrology. Malar J 12, 65

UNDP (2024). Technical Committee on Health “Mapping Risk for Vector-Borne Diseases (ID-V Risk)” Project Detection of Aedes aegypti in the island of Cyprus | United Nations Development Programme (undp.org), [https://www.undp.org/cyprus/publications/technical-committee-health-mapping-risk-vector-borne-diseases-id-v-risk-project-detection-aedes-aegypti-island-cyprus], Accessed 28/02/2024.

Vasquez M.I. et al. (2023). Two invasions at once: update on the introduction of the invasive species Aedes aegypti and Aedes albopictus in Cyprus - a call for action in Europe. Parasite, 2023; 30:41.

Violaris M, Vasquez MI, Samanidou A, Wirth MC, Hadjivassilis A. (2009). The Mosquito Fauna of the Republic of Cyprus: A Revised List. J. Am. Mosq. Control Assoc., 25:199–202.

Wint W et al. (2022). Past, present and future distribution of the yellow fever mosquito Aedes aegypti: The European paradox. Sci. Total Environ. 15; 847:157566.

World Bank (2021). World Integrated Trade Solution (WITS), Cyprus Food Products Imports & Exports by country in US$ Thousand 2021. Available at [https://wits.worldbank.org/CountryProfile/en/Country/CYP/Year/LTST/TradeFlow/Import/Partner/by-country/Product/16-24_FoodProd] & [https://wits.worldbank.org/CountryProfile/en/Country/CYP/Year/2021/TradeFlow/Export/Partner/by-country/Product/16-24_FoodProd], Accessed 28/02/2024.

