## Supplementary Materials for "The invasions of *Aedes aegypti* and *Aedes albopictus* in Cyprus: current situation, risk modelling and public health implications for the wider Eastern Mediterranean region"

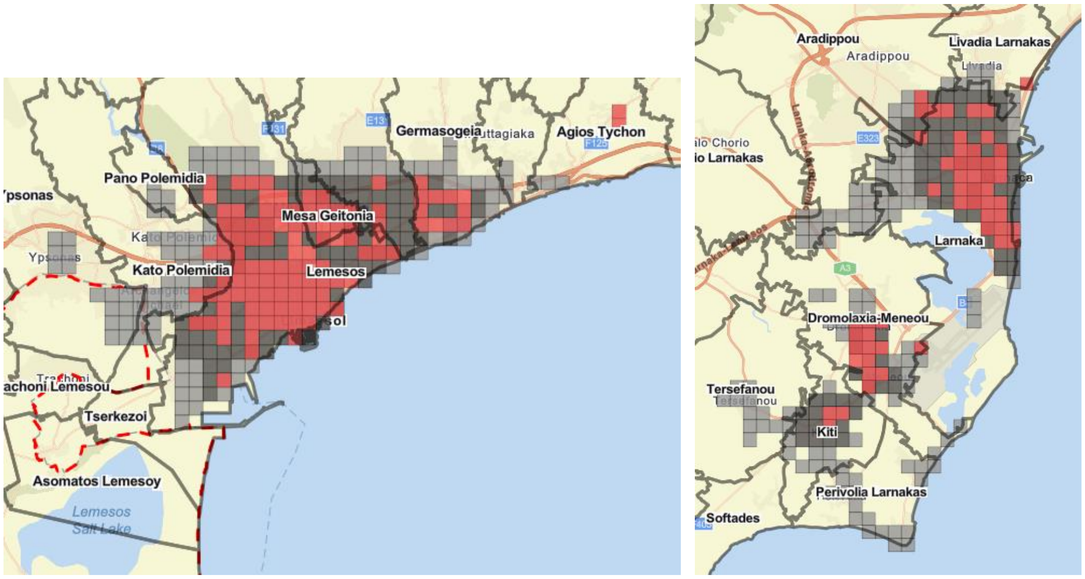


**Fig S1:** Mapping of 500 x 500m cells used for the delimitation sampling strategy of *Ae. albopictus* in Limassol (left) and *Ae. aegypti* in Larnaca (right).

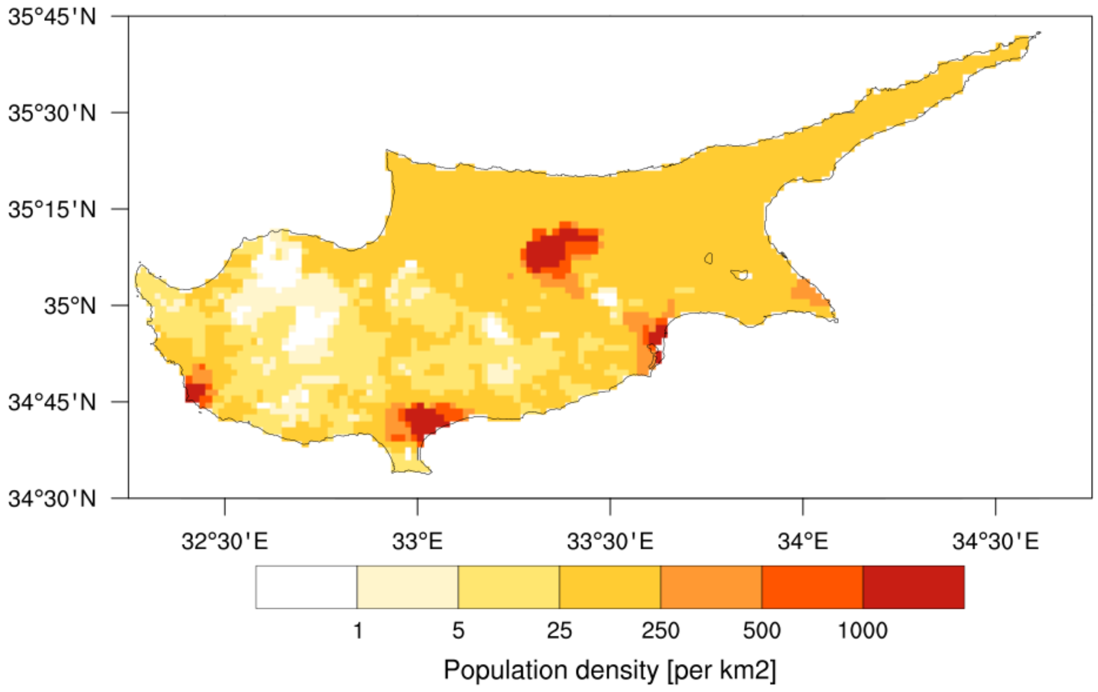


**Fig S2:** Population density (per km$^2$) for 2015 (UN adjusted data).

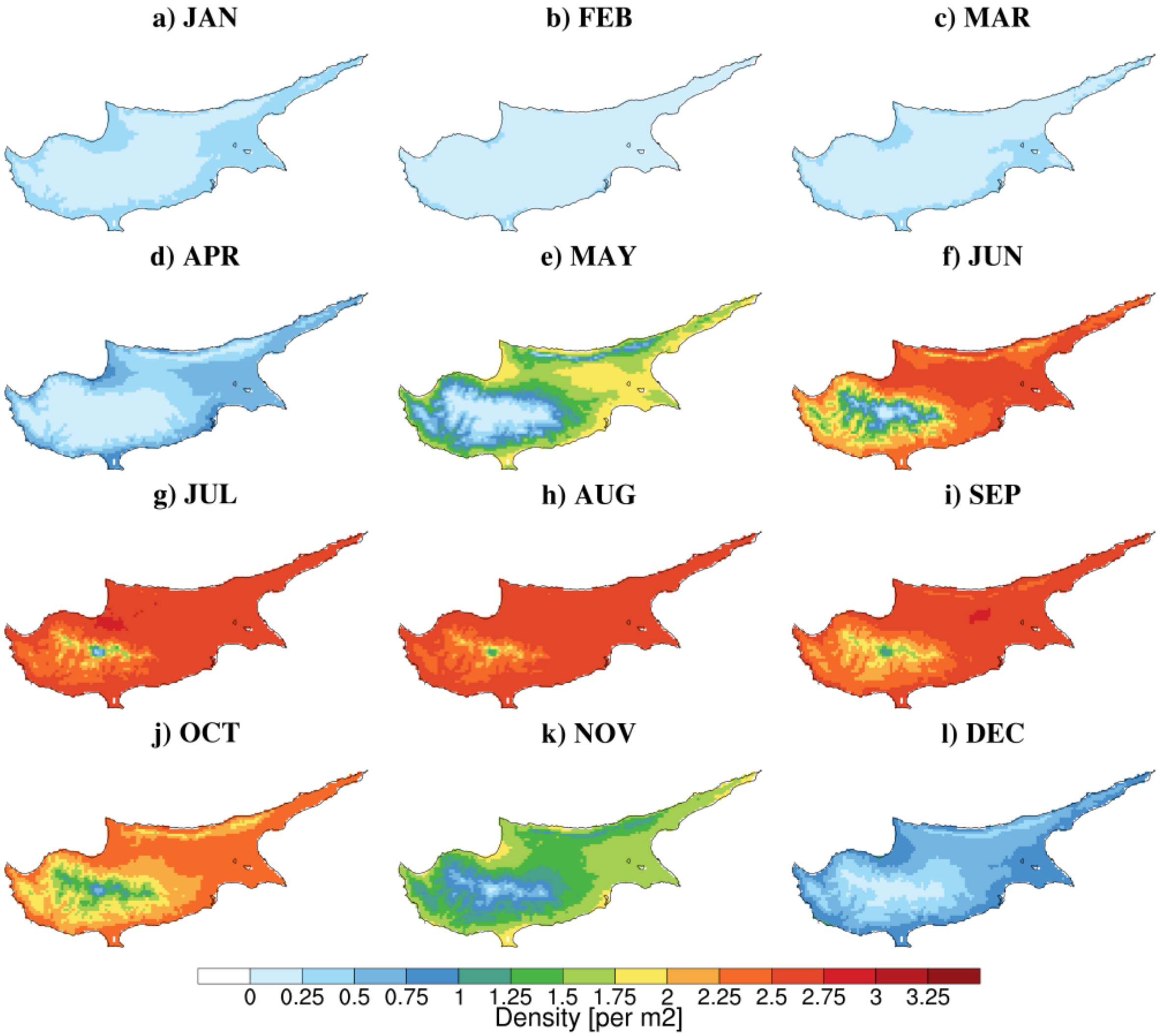


**Fig S3:** Mean seasonal cycle of simulated larval density for *Ae. aegypti* (1990-2022).

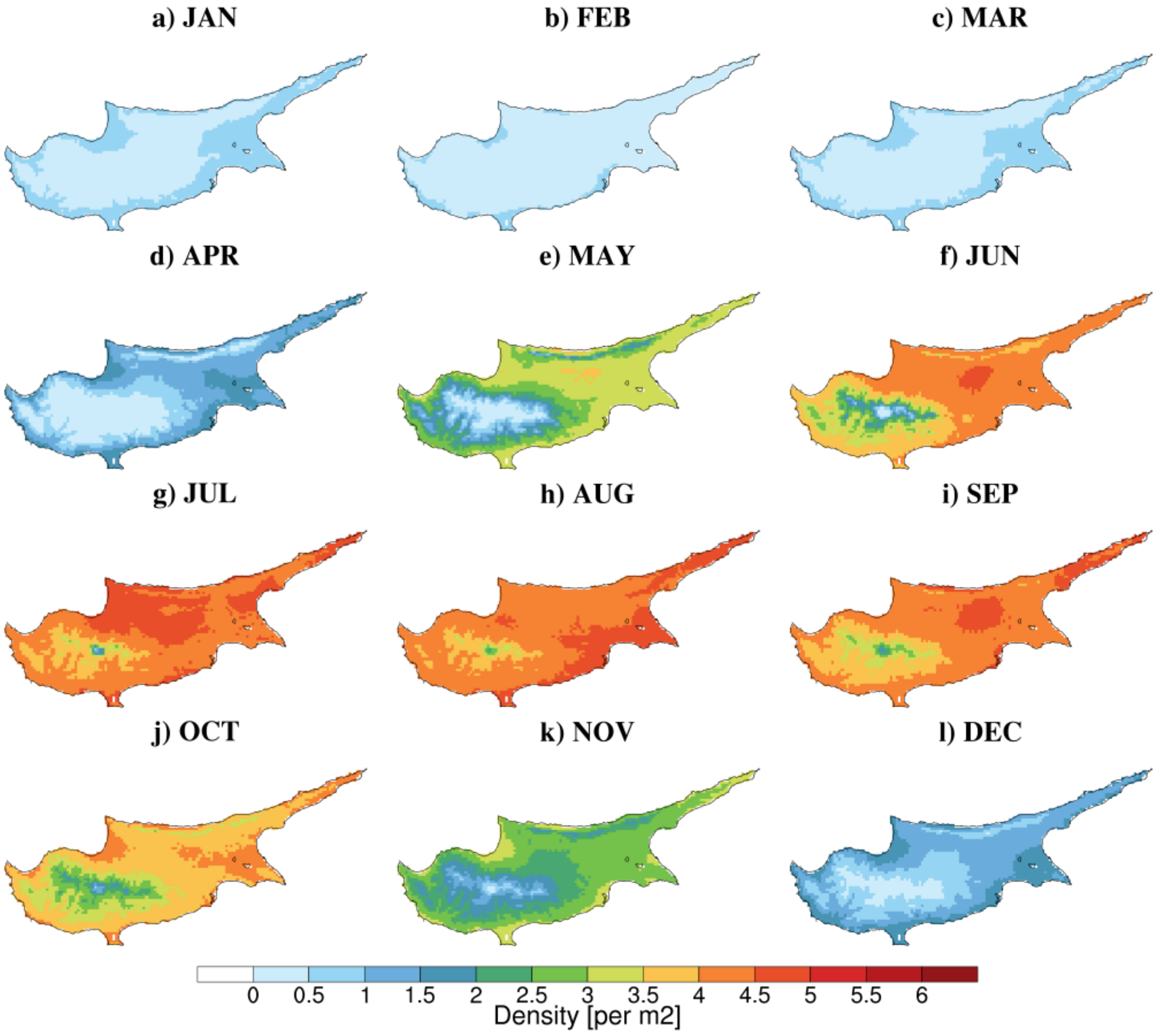


**Fig S4:** Mean seasonal cycle of simulated larval density for *Ae. albopictus* (1990-2022).

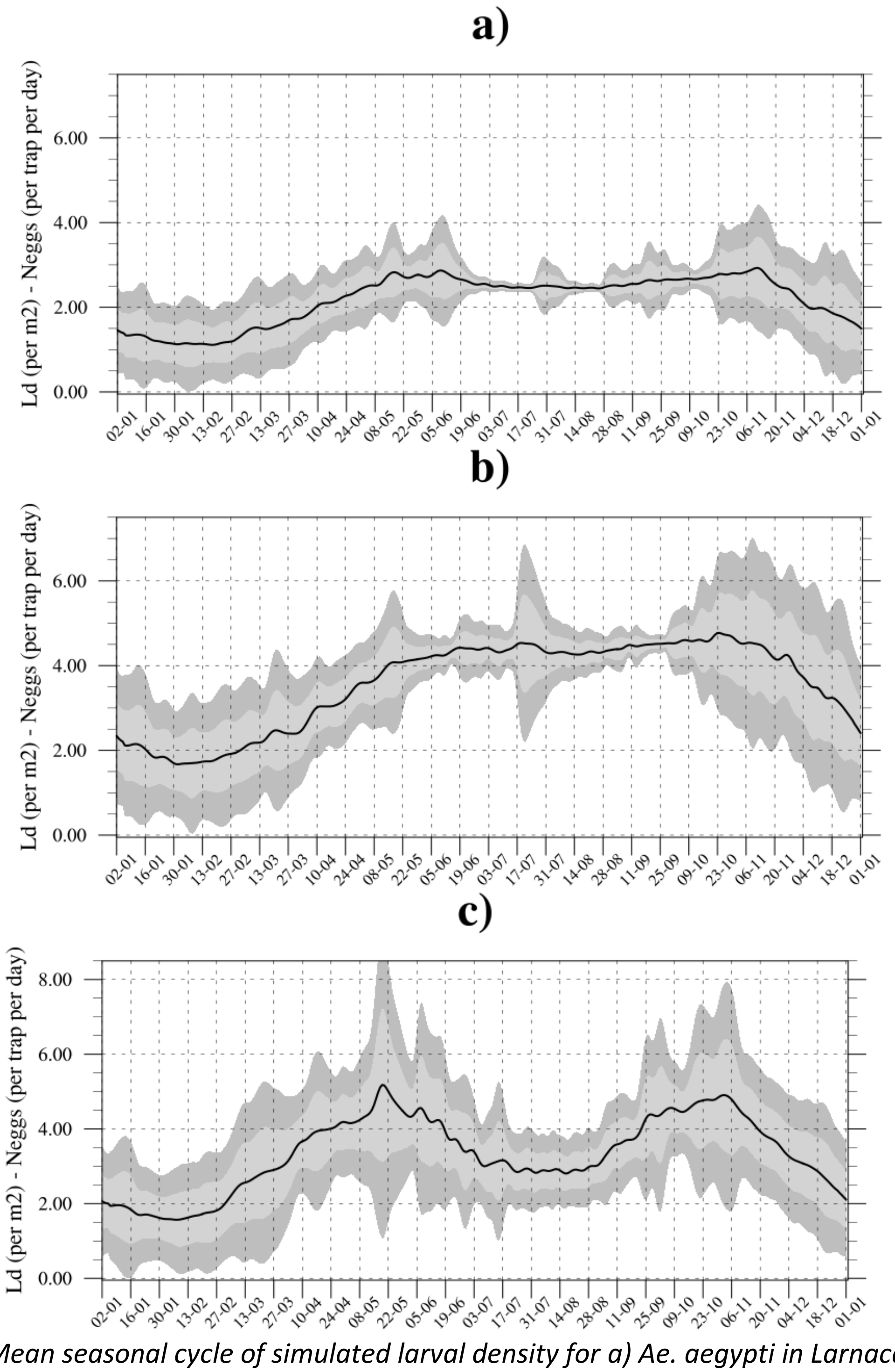


***Fig. S5****: Mean seasonal cycle of simulated larval density for a) Ae. aegypti in Larnaca, b) Ae. albopictus in Limassol and c) Nicosia. The mean for 1990-2022 is depicted by the black solid line. The light-gray and gray envelope respectively show one and two standard deviations around the mean calculated for all years. The VECTRI model was driven by daily maximum temperatures instead of mean temperatures.*
